# Cytoadhesion of *Plasmodium falciparum*-infected red blood cells changes the expression of cytokine-, histone- and antiviral protein-encoding genes in brain endothelial cells

**DOI:** 10.1101/2024.04.16.589757

**Authors:** Johannes Allweier, Michael Bartels, Hanifeh Torabi, Maria del Pilar Martinez Tauler, Nahla Galal Metwally, Thomas Roeder, Thomas Gutsmann, Iris Bruchhaus

## Abstract

Malaria remains a significant global health problem, mainly due to *Plasmodium falciparum*, which is responsible for most fatal infections. Infected red blood cells (iRBCs) evade spleen clearance by adhering to endothelial cells (ECs), triggering capillary blockage, inflammation, endothelial dysfunction and altered vascular permeability, prompting an endothelial transcriptional response. The iRBC^IT4var04^/HBEC-5i model, where iRBCs present IT4var04 (VAR2CSA) on their surface, was used to analyse the effects of iRBC binding on ECs at different temperature (37°C vs. 40°C). Binding of non-infected RBCs (niRBCs) and fever alone altered the expression of hundreds of genes in ECs. Comparing the expression profile of HBEC-5i cells cultured either in the presence of iRBCs or in the presence of niRBCs revealed significant upregulation of genes linked to immune response, nucleosome assembly, NF-kappa B signaling, angiogenesis, and antiviral immune response/interferon-alpha/beta signaling. Raising the temperature to 40°C, simulating fever, led to further upregulation of many genes, particularly those involved in cytokine production and angiogenesis. In summary, the presence of iRBCs stimulates ECs, activating several immunological pathways and affecting antiviral (- parasitic) mechanisms and angiogenesis. Our data uncovered the induction of the interferon-alpha/beta signaling pathway in ECs in response to iRBCs.

## 1 Introduction

Despite recent success stories in vaccine development and roll-outs, malaria remains one of the world’s most dangerous infectious diseases. In 2022, there were approximately 249 million cases of malaria, including 608,000 deaths, with most fatal malaria infections caused by *Plasmodium falciparum* (WHO, 2023). A vital aspect of the pathogenesis of *P. falciparum* infection is the cytoadhesion of infected red blood cells (iRBCs) to the vascular bed of critical organs such as the brain, heart, lungs, stomach, skin, and kidneys (D. Milner, Jr. et al., 2013; D. A. Milner, Jr. et al., 2015; Taylor et al., 2004). In addition to capillary blockage, iRBC cytoadhesion induces inflammatory cytokine production, endothelial dysfunction, and vascular permeability in the affected tissue (Lyke et al., 2004). As a result of the immune responses triggered by parasite growth and endothelial cytoadhesion, patients may develop fever, headache, myalgia, and muscle stiffness (Cunnington et al., 2013; Hasday et al., 2001; Oakley et al., 2011; Oakley et al., 2007; Phillips et al., 2017). Depending on the age and immune status of the patient, severe to life-threatening complications may develop, including cerebral malaria, organ failure, acidosis, and severe anemia (Cunnington et al., 2013; Gazzinelli et al., 2014; Phillips et al., 2017).

Cytoadhesion is primarily mediated by members of the *P. falciparum* erythrocyte membrane protein 1 (*Pf*EMP1) family, encoded by approximately 45-90 *var* genes per parasite genome (Otto et al., 2019). The expression of *var* genes is mutually exclusive in ring-stage parasites, so only one *Pf*EMP1 variant is present on the surface of trophozoite- and schizont-stage iRBCs at any one time. The *var* genes encoding *Pf*EMP1 are highly variable from parasite to parasite, resulting in a vast repertoire of *var* genes in nature (Deitsch & Dzikowski, 2017; Kraemer & Smith, 2006; Kyes et al., 2007; Pasternak & Dzikowski, 2009; Voss et al., 2006). *Pf*EMP1s are concentrated in electron-dense protrusions of the erythrocyte plasma membrane called knobs (Sanchez et al., 2019). Knobs are composed of several submembrane structural proteins, including the major protein of this structure, the knob-associated histidine-rich protein (KAHRP)(Alampalli et al., 2018; Gruenberg et al., 1983; Jager et al., 2022; Maier et al., 2009; Sanchez et al., 2022; Tilly et al., 2015) and play an essential role in binding iRBCs to the endothelium (Crabb et al., 1997; Cutts et al., 2017; Dorpinghaus et al., 2020; Horrocks et al., 2005; Lubiana et al., 2020). If iRBCs do not form knobs, e.g. due to loss of genes encoding knob structural proteins, adhesion under flow conditions is significantly impaired. Especially at febrile temperatures, they play a crucial role by stabilizing the binding interaction. Under these conditions, knob-negative iRBCs can no longer bind to various endothelial cell receptors (ECRs) (Lubiana et al., 2020).

Cytoadhesion of iRBCs also changes gene expression in endothelial cells (ECs) (Tripathi et al., 2009; Tripathi et al., 2006). Stimulation of primary human brain microvascular endothelial cells (HBMECs) through cytoadhesion of iRBCs resulted in increased ICAM-1 expression on the surface of HBMECs, resulting in increased cytoadhesion. This observation was specific to HBMECs, in contrast to human umbilical vein endothelial cells (HUVECs), where no significant changes in gene expression were observed (Tripathi et al., 2006). In another study, the influence of cytoadhesion on HBMEC gene expression was investigated by microarray analysis (Tripathi et al., 2009). The effect of binding of iRBCs with enhanced binding on HBMECs induced by panning was compared with the effects of binding of unpanned iRBCs. For both populations, a 4-fold increase of ICAM-1 presented on the HBMEC surface was detected after 6 h of co-incubation. Interestingly, there were no significant differences in gene expression between the two parasite populations. Using GO term analysis, the proteins encoded by the regulated genes were shown to be involved in inflammatory response, apoptosis, anti-apoptosis, cell-cell signaling and transduction, and NF-κB activation cascade (Tripathi et al., 2009). Recently, the transcriptome profile of HBMECs co-cultured for different time points with iRBCs presenting IT4_var14 (strong binding to ICAM-1) or IT4_var37 (strong binding to CD36) on their surface was analysed. Genes coding for proteins involved in many of the pathways described above, such as inflammation, cell-cell interaction, and apoptosis, were identified as differentially expressed. However, significant differences were also found between iRBC^IT4var14^ and iRBC^IT4var37^. Among others, atonal bHLH transcription factor 8 (*ATOH8*), *CXCL8*, *CXCL10*, *VCAM-1*, radical s-adenosyl methionine domain containing 2 (*RSAD2*), and *IL6* were found to be up-regulated in the presence of iRBC^IT4var14^ after 6 h of co-cultivation. In contrast, whereas the expression of these genes was down-regulated in iRBC^IT4var37^ (Othman et al., 2023).

Chakravorty and colleagues also investigated the influence of cytoadhesion on HUVEC gene expression (Chakravorty et al., 2007). They confirmed the observation of Tripathi and colleagues that cytoadhesion of iRBCs alone is insufficient to stimulate HUVEC cells (Tripathi et al., 2006). However, when co-incubated in the presence of 5 pg ml^−1^ of TNF, there is an increase in the presentation of ICAM-1 on the cell membrane. Using microarray analysis, they showed that genes encoding proteins involved in cell communication, cell adhesion, signal transduction, and immune response were regulated. Thus, HUVEC can mobilize immune and adhesion reactions, but only when co-incubated with low concentrations of TNF in addition to iRBCs cytoadhesion (Chakravorty et al., 2007).

In previous studies, an immortalized human cerebral microvascular endothelial cell line (HBEC-5i) was used to characterize the cytoadhesion of iRBCs (Dorpinghaus et al., 2020; Lubiana et al., 2020). HBEC-5i cells are derived from the microvasculature of the human cerebral cortex and, therefore, have essential characteristics of cerebral ECs. HBEC-5i cells present Willebrand factor (vWF), VE-cadherin, occludin, VCAM-1, ICAM-1, CD54, CD40, and chondroitin-sulfate A (CSA) on the cell surface. However, the binding of iRBCs of the *P. falciparum* isolate IT4 to HBEC-5i cells was shown to occur mainly via CSA. Parasites enriched for binding to HBEC-5i cells present IT4var04 (VAR2CSA) on the surface of iRBCs (iRBCs^IT4var04^), of which CSA is known to be the ligand. Thus, CSA seems to be dominant on HBEC-5i cells, and the binding of iRBCs to other receptors, such as ICAM-1, is shielded (Dorpinghaus et al., 2020). This model (iRBC^IT4var04^/HBEC-5i) showed that binding to CSA occurs only at a low shear stress of 0.9 dyne/cm^2^ with a static binding characteristic. Additionally, it could be shown that binding at febrile temperatures can only occur when the iRBCs have formed knobs (Lubiana et al., 2020).

In the present study, the iRBC^IT4var04^/HBEC-5i model was used to investigate whether stimulation of ECs by binding of iRBCs to CSA is possible and, if so, whether and how this stimulation differs from that of HBMEC and HUVEC and what happens in the presence of fever. The aim was to determine whether cytoadhesion, in general, elicits similar responses in ECs, independent of the receptor. Moreover, we also studied the effects of febrile temperatures due to the co-incubation with iRBCs.

**2 Results**

### 2.1 Effects of niRBCs, iRBCs, and fever on the expression of HBEC-5i cells

By performing a complete transcriptome analysis of the ECs, we examined the impact of niRBCs, iRBCs, and fever on the expression profile of HEBEC-5i cells. For these studies, as mentioned above, the *P. falciparum* isolate IT4 was used, which were enriched on HBEC-5i cells over several rounds, with adhesion taking place via CSA, thus presenting IT4var04 (VAR2CSA) on the surface of the iRBCs (Dorpinghaus et al., 2020). A Venn diagram shows the differential expression (true vs. false) of the three parameters niRBC, iRBC, and febrile (Figure 1). Each circle represents the differential expression (true vs. false) for the three parameters. Intersections represent genes that are differentially expressed in two or three analyses. Genes exhibiting a differential expression of a fold change of ≥ 2.0, a false discovery rate (FDR) of ≤ 0.05, and a minimum transcript per million (TPM) threshold of ≥ 2.0 were included in this analysis.

**Figure 1.**
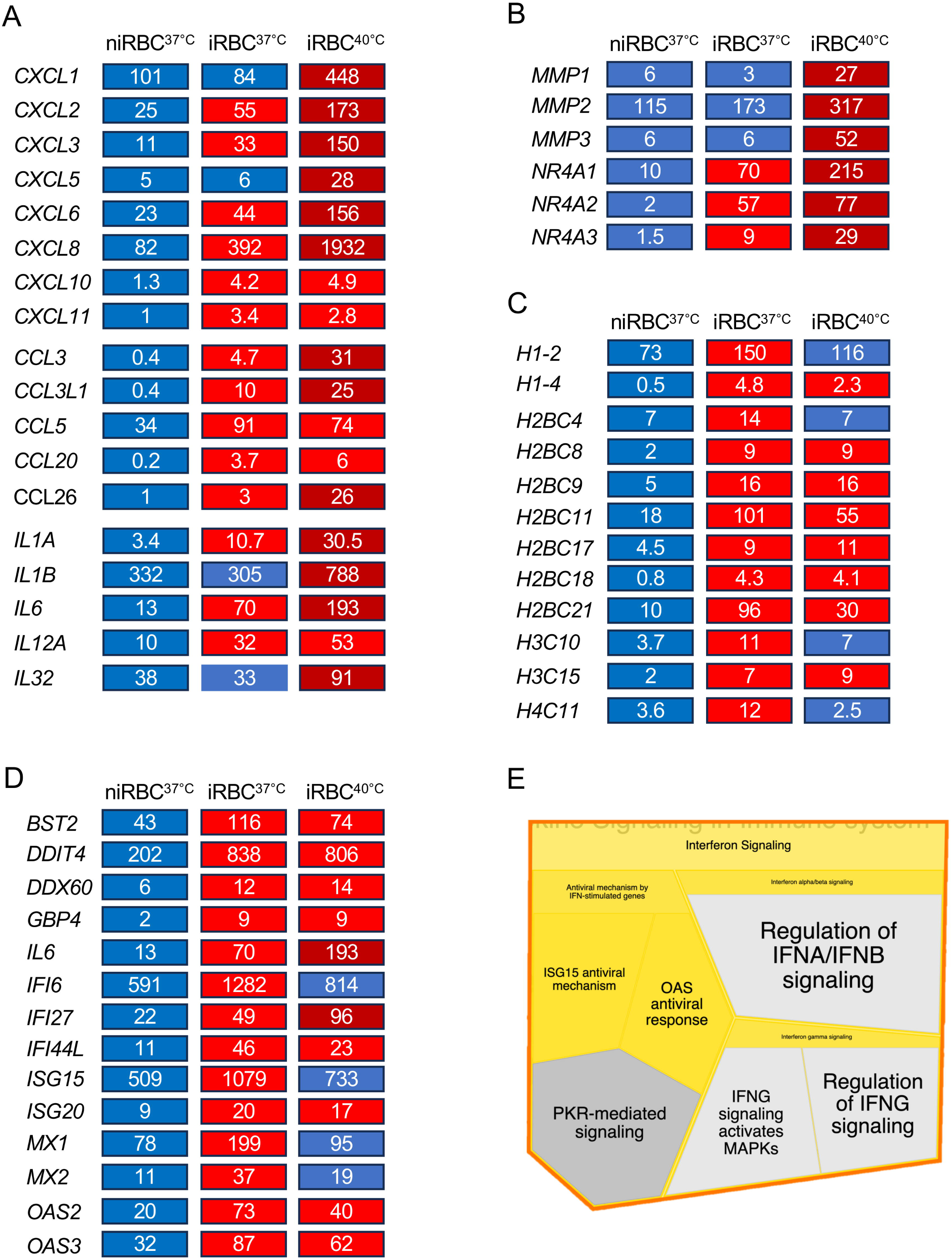
Venn diagram showing the number of overlapping and differentially expressed genes in HBEC-5i cells of the groups niRBC true vs. false (yellow circle), febrile true vs. false (red circle) and iRBC true vs. false (blue circle). The niRBC true vs. false group contains the genes for which the presence of niRBC affects the expression of HBEC-5i cells. The febrile true vs. false group contains genes for which increasing the culture temperature from 37°C to 40°C has an effect on the expression of HBEC-5i cells. The group iRBC true vs. false contains the genes for which the presence of iRBCs changes the expression of HBEC-5i cells. Each circle represents one of the groups, with overlaps including genes present in more than one group. Fold change ≥ 2.0, false discovery rate (FDR) ≤ 0.05, minimum TPM threshold ≥ 2.0.

Our study has provided clear results, revealing distinct regulatory effects: 60 genes were exclusively modulated by the presence of niRBCs, while a mere increase intemperature from 37°C to 40°C led to altered expression patterns in 98 genes.. Similarly, the presence of iRBCs alone influenced the expression of 318 genes. Notably, 285 genes exhibited regulation in response to either niRBCs or iRBCs, with 80 genes showing such regulation specifically in the presence of iRBCs^37°C^ and iRBCs^40°C^. Additionally, ten genes were differentially expressed either by the febrile stimulus or presences of niRBCs. Seventy-nine genes were found that overlap in all three comparisons (Figure 1, Supplementary Table 1).

Of the 59 genes significantly differentially expressed by niRBCs, 38 are up-regulated and 21 are down-regulated. With minor exceptions, most of these genes demonstrate a 2- to 3-fold regulation. Notably, the most highly up-regulated gene, BOLA2, shows a striking 79-fold increase in response to the presence of niRBCs (from TPM 2.5 to 198.3) (Figure 1, Supplementary Table 1).

Among the 98 genes significantly differentially expressed by the temperature elevation from 37°C to 40°C, 50 show up-regulation, while 48 undergo down-regulation. Most of these genes demonstrate a 2- to 3-fold regulation. Notably, the increase in temperature triggers the up-regulation of 10 genes associated with the Gene Ontology Biological Process (GO-BP) term “protein folding” (FDR 4.82e-16) (Figure 1, Figure 2A, Supplementary Table 1). This observation highlights the biological response to the environmental change of temperature. Analysing the 318 genes significantly differentially expressed by the presence of iRBCs, 121 are up-regulated, while 197 are down-regulated (Supplementary Table 1). Approximately 90% of these genes undergo a 2- to 3-fold regulation, while the remaining 10% (22 genes) exhibit a 3- to 6-fold regulation. String analysis reveals the up-regulation of GO-BP terms such as “defense response to virus” (FDR 7.75e-09), “nucleosome assembly” (FDR 0.00013), “NF-κB complex” (FDR 0.0018), and “ribosome biogenesis” (FDR 0.00086) (Figure 1, Figure 2B, Supplementary Table 1).

**Figure 2.**
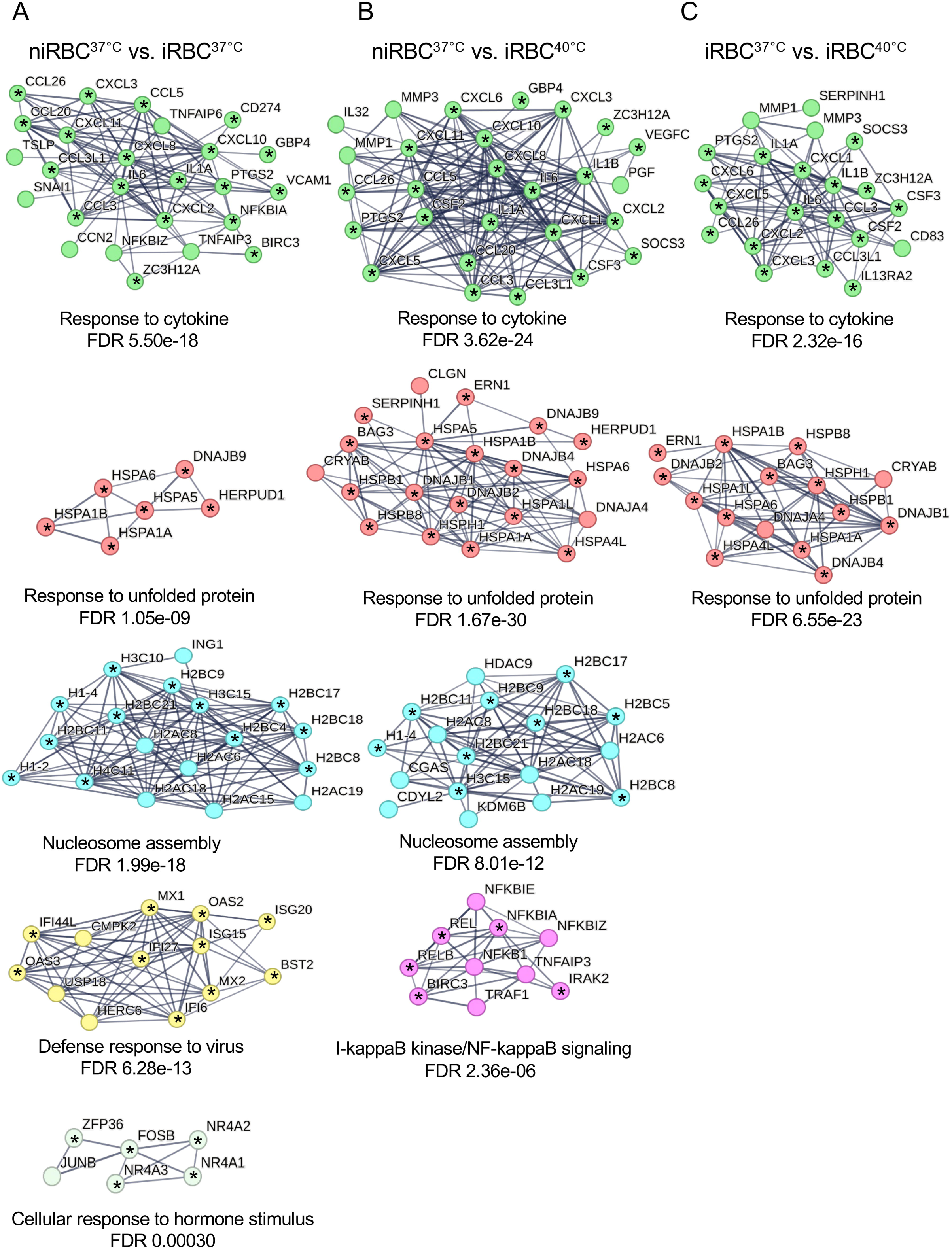
Protein-protein interaction network analysis. A STRING analysis was performed to identify protein-protein interaction networks that are regulated in HBEC-5i cells in response to the presence of niRBCs, febrile temperature and/or iRBCs. The nodes represent proteins and the connecting lines between the nodes indicate known or predicted interactions. All interaction networks that are shown are up-regulated compared to the respective controls. The thickness of the connecting lines indicates the confidence level of the interaction. Nodes marked with an asterisk are proteins whose encoding genes were identified as significantly differentially expressed in the respective analyses. A selection of significant networks from the GO-BP analysis is shown. (**A**) Comparison febrile true vs. false, contains 98 group-specific significantly regulated genes. (**B**) Comparison iRBC true vs. false, contains 318 group-specific significantly regulated genes. (**C**) Intersection of the groups niRBC true vs. false and iRBC true vs. false. (**D**) Intersection of the groups iRBC true vs. false and febrile true vs. false. (**E**) Intersection of the groups niRBC true vs. false, iRBC true vs. false, and febrile true vs. false. Genes exhibiting significant differential expression, meeting stringent criteria including a fold change of ≥ 2.0, a FDR of ≤ 0.05, and a minimum threshold of TPM of ≥ 2.0, were considered for inclusion in the analysis.

Within the intersection of genes derived from the analyses of niRBC true vs. false and iRBCs true vs. false, a total of 285 genes exhibit regulation. Noteworthy among these are 13 genes associated with “cellular response to cytokine stimulus” (FDR 4.82e-10), 9 genes linked to

“nucleosome assembly” (FDR 2.56e-15), 11 genes involved in “I-κB kinase/NF-κB signaling” (FDR 4.32e-06), 5 genes contributing to “negative regulation of I-κB kinase/NF-κB signaling” (FDR 0.0017), and 3 genes associated with “defense response to virus” (FDR 3.38e-05) GO-BP terms. Remarkably, all these pathways demonstrate up-regulation (Figure 1, Figure 2C, Supplementary Table 1).

Within the intersection encompassing genes regulated in both the iRBC true vs. false and febrile true vs. false analyses, a cohort of 80 genes is observed. Among this group, three genes are associated with the “vascular endothelial growth factor signaling pathway” GO-BP term (FDR 0.00025). In comparison, four genes are linked to the “response to unfolded protein” term (FDR 2.50e-05), all demonstrating up-regulation (Figure 1, Figure 2D, Supplementary Table 1).

In the intersection of all three comparative analyses (niRBC true vs. false; febrile true vs. false; iRBC true vs. false), a set of 80 regulated genes is identified. Notably, the pathways significantly up-regulated include the “chemokine-mediated signaling pathway” (FDR 2.26e-12), comprising 13 genes, and the “protein folding” GO-BP term (FDR 3.29e-10), comprising seven genes (Figure 1, Figure 2E, Supplementary Table 1).

In the analysis “iRBCs true vs. false”, which comprises 762 genes, a comprehensive string analysis reveals 7 distinct clusters. These clusters house 6 to 24 of the regulated genes and correspond to the GO-BP terms “response to cytokine” (FDR 2.01e-23; 24 genes), “response to unfolded protein” (FDR 1.67e-30; 17 genes), “positive regulation of RNA metabolic process” (FDR 7.04e-10; 15 genes), “nucleosome organization” (FDR 3.83e-11; 9 genes), “defense response to virus” (FDR 7.04e-16; 12 genes), and “I-κB kinase/NF-κB signaling” (FDR 3.24e-08; 6 genes). Remarkably, all these clusters exhibit up-regulation except for the sole down-regulated cluster pertaining to “nucleosome assembly” (FDR 3.64e-08; 7 genes). These detailed findings provide a comprehensive understanding of the gene regulations in the ‘iRBCs true vs. false’ analysis (Figure 1, Figure 3A, Supplementary Table 1).

**Figure 3.**
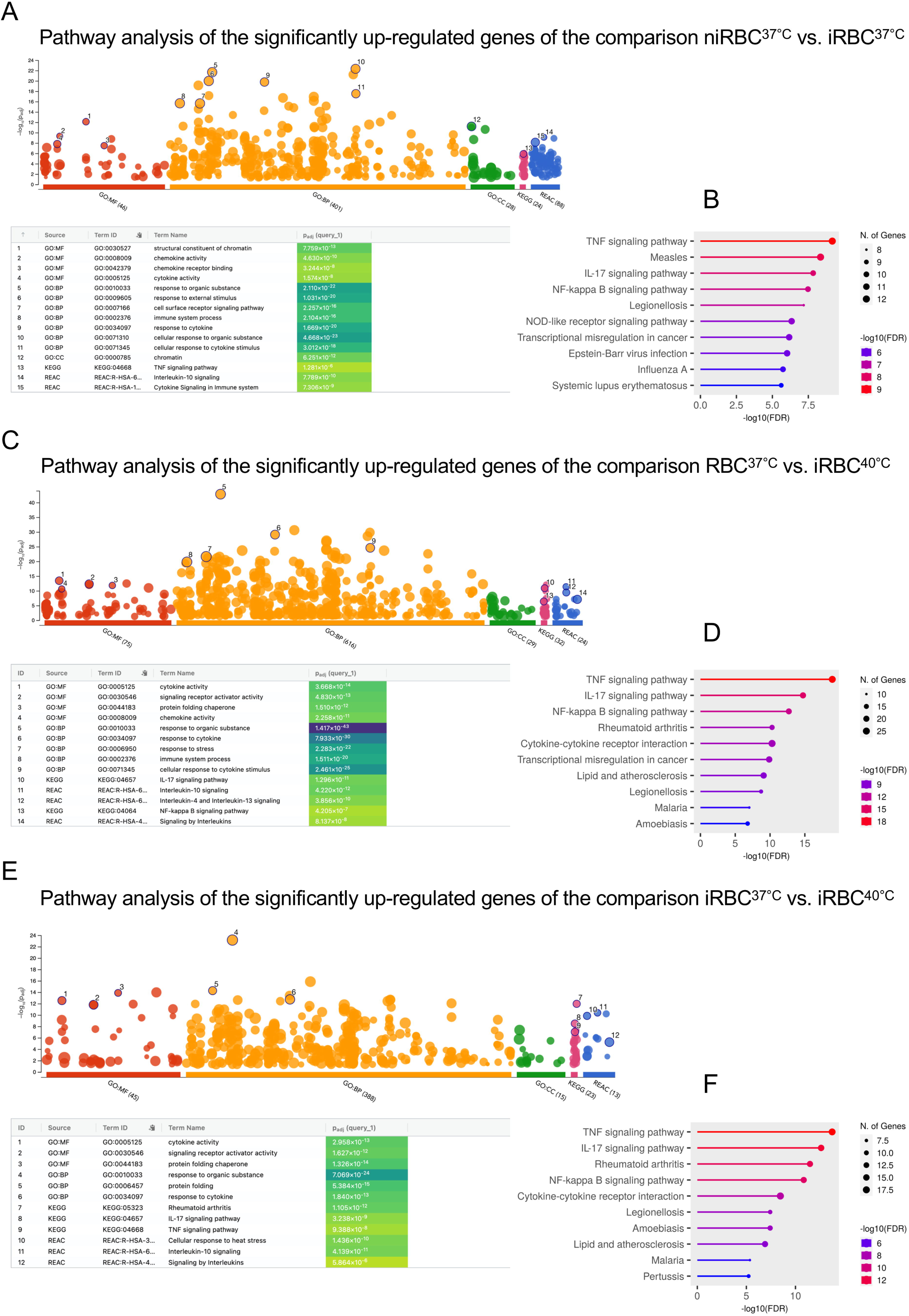
Identification of the pathways regulated by the presence of iRBCs in HBEC-5i cells. (**A**) A STRING analysis was performed to identify protein-protein interaction networks regulated in HBEC-5i cells in response to the presence of iRBCs (iRBC true vs. false including 762 genes). The nodes represent proteins and the connecting lines between the nodes indicate known or predicted interactions. The thickness of the connecting lines indicates the confidence level of the interaction. Nodes marked with an asterisk are proteins whose encoding genes were identified as significantly differentially expressed in the analyses. A selection of significant networks from the GO-BP analysis is shown. Genes exhibiting significant differential expression, meeting stringent criteria including a fold change of ≥ 2.0, a FDR of ≤ 0.05, and a minimum threshold of TPM of ≥ 2.0, were considered for inclusion in the analysis. (**B**) ShinyGO analysis to categorize the proteins encoded by the identified genes according to their biological functions. The 10 most significant pathways are shown.

Using ShinyGO analysis, the proteins encoded by the identified genes were categorized according to their biological functions. Among the ten most significantly regulated functions are the TNF, IL-17, and NF-κB signaling pathways and cytokine-cytokine receptor interaction, as well as the response to viral (herpes simplex virus 1 infection) and bacterial (legionellosis) infections (Figure 3B).

### 2.2 Specific response of HBEC-5i cells to cytoadhesion of iRBCs^37°C^ and iRBCs^40°C^

It is important to emphasize that in the above comparisons, although the expression of genes within different intersections is altered compared to the respective controls, the magnitude of these changes in expression levels is not considered in the analyses. Hence, a more detailed investigation is warranted to delineate the specific impact of iRBC cytoadhesion on HBEC-5i cells and the role of the cultivation temperature (37°C vs. 40°C). In contrast to the previous analysis, this subsequent investigation only includes the mean values of normalized reads from three biologically independent transcriptomes. However, the focus remains exclusively on genes identified as significantly regulated within the analysis above (Supplementary Table 1). In the first step, we identified those genes whose expression was up- or down-regulated (fold change ≥ 2.0, FDR ≤ 0.05, TPM threshold ≥ 2.0) in HBEC-5i cells in response to iRBCs compared to the presence of niRBCs at 37°C (iRBCs^37°C^ vs. niRBCs^37°C^). Cytoadhesion of iRBCs^37°C^ up-regulates the expression of 162 genes and down-regulates the expression of 219 genes compared to the presence of niRBCs^37°C^ (Supplementary Table 1).

To get an idea of which metabolic pathways the proteins encoded by the 162 up-regulated genes are involved, a g:Profiler analysis was performed. The GO Molecular Function (GO-MF) terms “structural component of chromatin” and “chemokine activity” had the highest significance. For the GO-BP terms, these were “cellular response to organic substance” and “response to cytokine”. The KEGG pathway analysis showed the highest significance for “TNF signaling pathway”, and in the Reactome analysis these were “interleukin-10 signaling” and “cytokine signaling immune system” (Figure 4A). The ShinyGO analysis shows similar results. Here, as described in the previous paragraph, different signaling pathways of the immune system and the response to viral and bacterial infections were mainly regulated (“TNF, IL-17 and NF-κB signaling pathways”, “measles”, “legionellosis”) (Figure 4B). The use of a string analysis yielded very similar results. Here, among the 162 up-regulated genes, the KEGG pathways exhibiting the most stringent regulation include the “TNF signaling pathway” (FDR 6.70e-09), “viral protein interaction with cytokine and cytokine receptor” (FDR 1.03e-07), “IL-17 signaling pathway” (FDR 4.42e-07), “cytokine-cytokine receptor interaction” (FDR 7.64e-07), and “NF-κB signaling pathway” (FDR 8.31e-07). In total, eight genes are up-regulated more than 10-fold, while another 16 genes are up-regulated more than 5-fold. Notable among these are the genes encoding the chemokines CCL31, CCL3, CCL20, and IL-6 (Supplementary Table 1).

**Figure 4.**
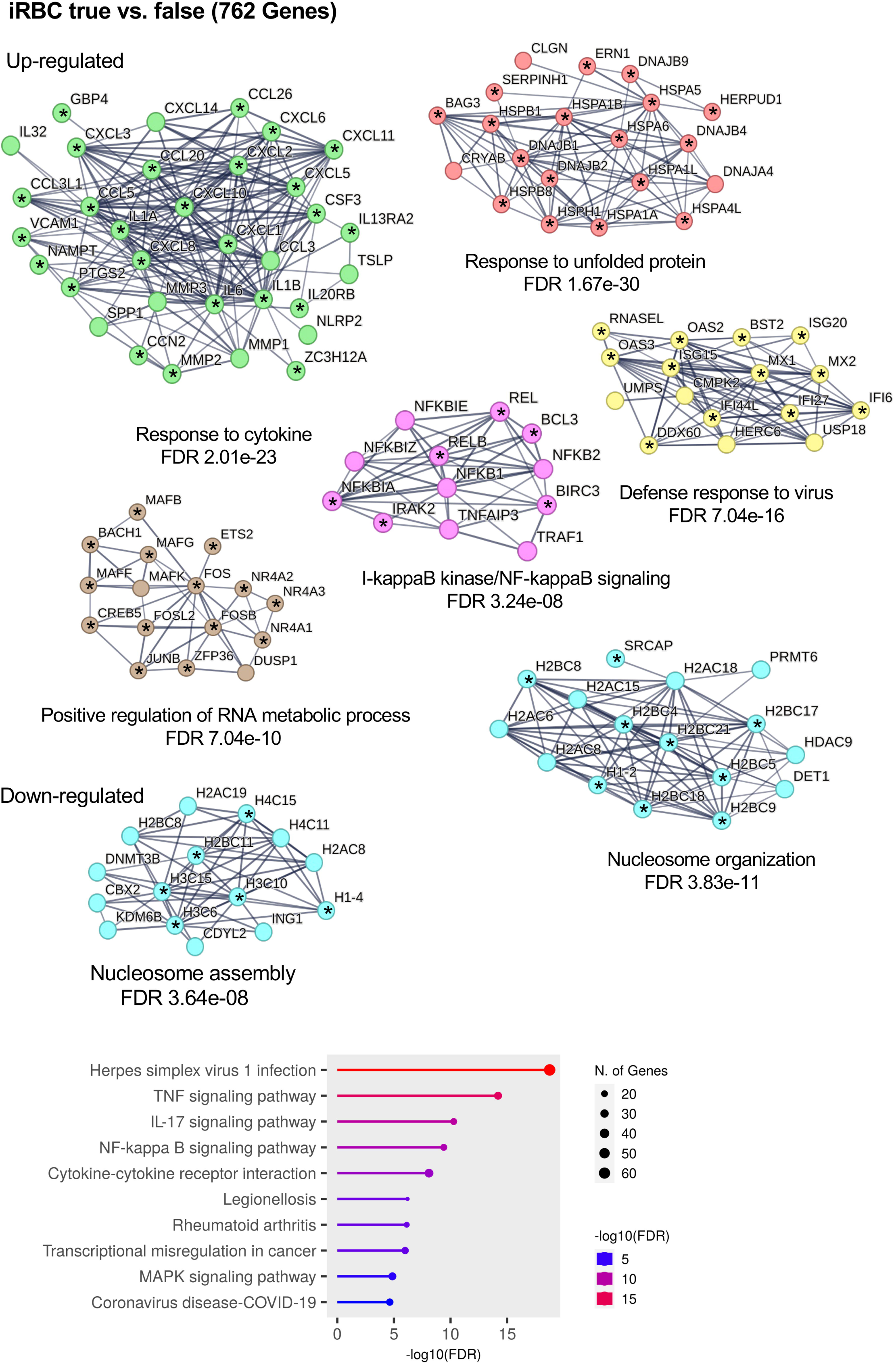
Pathway analysis of the specific response of HBEC-5i cells due to the cytoadhesion of iRBCs^37°C^ and iRBCs^40°C^ co-cultivation. g:Profiler ((**A**), (**C**), (**E**)) and ShinyGO ((**B**), (**D**), (**F**)) analysis of the significantly up-regulated genes (fold change ≥ 2.0, FDR ≤ 0.05, and minimum threshold of TPM of ≥ 2.0) of the comparisons (**A**), (**B**): niRBC^37°C^ vs. iRBC^37°C^; (**C**), (**D**): niRBC^37°C^ vs. iRBC^40°C^; (**E**), (**F**): iRBC^37°C^ vs. iRBC^40°C^.

Furthermore, the up-regulated genes cluster into the following GO-BP terms: “response to cytokine” (FDR 5.50e-18, 19 genes), “nucleosome assembly” (FDR 1.99e-18, 12 genes), “defense response to virus” (FDR 6.28e-13, 10 genes), “cellular response to hormone stimulus” (FDR 0.00030, 5 genes). (Figure 5A, Supplementary Table 1). The 219 genes whose expression is down-regulated code for proteins that fall under the GO-MF term “DNA-binding transcription factor activity” (FDR 2.008×10^−20^) and the GO-BP term “regulation of nucleobase-containing compound metabolic process” (FDR 3.664×10^−18^) according to the g:Profiler analysis (Supplementary Table 1).

**Figure 5.**
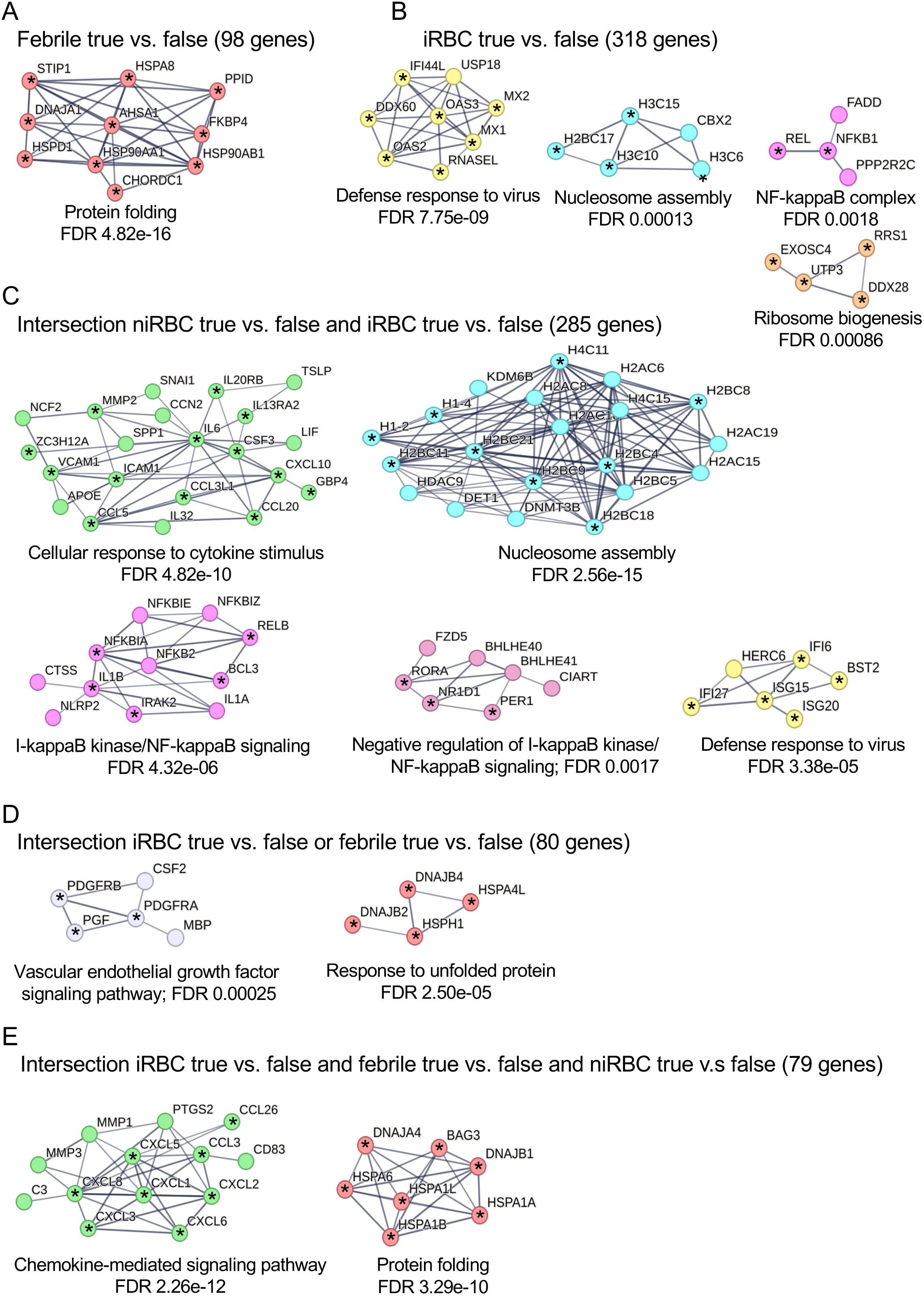
Identification of the pathways regulated by the presence of iRBCs in HBEC-5i cells at 37°C and 40°C using a STRING analysis. Nodes represent proteins and the connecting lines between the nodes indicate known or predicted interactions. The thickness of the connecting lines indicates the confidence level of the interaction. Nodes marked with an asterisk are proteins whose encoding genes were identified in the analyses. A selection of significant networks from the GO-BP analysis is shown. Genes exhibiting significant differential expression, meeting stringent criteria including a fold change of ≥ 2.0, a FDR of ≤ 0.05, and a minimum threshold of TPM of ≥ 2.0, were considered for inclusion in the analysis. STRING analysis was performed to identify protein-protein interaction networks regulated in HBEC-5i cells after co-cultivation of (**A**) iRBCs^37°C^ in comparison to niRBCs^37°C^, (**B**) iRBCs^40°C^ in comparison to niRBCs^40°C^, and **C** iRBCs^37°C^ in comparison to iRBCs^40°C^.

The next question was to investigate the effect of increasing the temperature from 37°C to 40°C (iRBCs^40°C^), simulating fever as an additional factor besides cytoadhesion of iRBCs^37°C^. Therefore, the expression profile of HBEC-5i cells following cytoadhesion iRBCs^40°C^ was compared to HBEC-5i cells cultured in the presence of niRBCs^37°C^. This analysis revealed 381 genes up-regulated and 213 genes down-regulated in expression (fold change ≥ 2.0, FDR ≤ 0.05, TPM threshold ≥ 2.0) (Supplementary Table 1). G:Profiler and ShinyGO analyses group of the 381 genes up-regulated at 40°C were also carried out here. By and large, the same pathways were identified. However, the significance was more than doubled for many of the pathways, e.g., for the TNF signaling pathway (Figures 4C, 4D). A string analysis of this comparison also shows excellent agreement with the abovementioned analyses. The same KEGG pathways are regulated at 40°C that were already addressed at 37°C but with one exception with higher significance (“TNF signaling pathway” (FDR 1. 54e-17), “IL-17 signaling pathway” (FDR 2.06e-12), “cytokine-cytokine receptor interaction” (FDR 8.20e-11) and “NF-κB signaling pathway” (FDR 8.00e-11), “viral protein interaction with cytokine and cytokine receptor” (FDR 6.70e-07)). GO-BP terms identified as significant in the iRBC^37°C^ with niRBCs^37°C^ comparison, such as “defence reaction to viruses” and “cellular reaction to hormone stimuli”, are no longer found in the iRBC^40°C^ with niRBCs^37°C^ comparison. Instead of twelve genes, the GO term “nucleosome assembly” (FDR 8.01e-12) still shows nine genes as significantly up-regulated in their expression. The significance for the GO-BP term “response to unfolded protein” increases from FDR 1.09e-09 (6 genes) to FDR 1.67e-30 (17 genes). This is also true for the GO-BP term “cytokine response” (FDR 3.62e-24), to which 23 genes can be assigned. Furthermore, the GO-BP term “I-κB kinase/NF-κB signaling” (FDR 2.36e-06, 5 genes) was identified as regulated (Figure 4B, Supplementary Table 1). In addition, significant up-regulation of gene expression was detected, with a notable subset showing ≥10-fold increase (32 genes including *CCL26, CCL20, CCL3L1, CXCL3, CXCL8,* and *Il6*) and a further 55 genes showing 5- to <10-fold increase when comparing niRBC^37°C^ vs. iRBC^40°C^ (Supplementary Table 1).

The 213 genes whose expression is down-regulated encode for proteins that fall under the GO-MF term “DNA-binding transcription factor activity” (FDR 1.805×10^−12^) and the GO-BP term “regulation of transcription by RNA polymerase II” (FDR 4.185×10^−9^) according to the g:Profiler analysis (Supplementary Table 1).

Our investigation proceeded to scrutinize the genes with regulated expression in the presence of iRBCs at 40°C, contrasting them with those at 37°C (fold change ≥ 2.0, FDR ≤ 0.05, TPM threshold ≥ 2.0). We identified 205 genes (18 genes fold change >10-, 25 genes fold change 5-10) where cytoadhesion at 40°C results in a further increase in expression compared to cytoadhesion at 37°C. We observed down-regulation in 38 genes (Supplementary Table 1). Notably, some genes exhibited a continuous increase in expression, starting from cultivation in the presence of niRBCs^37°C^ to iRBCs^37°C^ to iRBCs^40°C^. For instance, this trend is evident in the case of *CXCL8*, where the expression rises from 11.7 TPM (no niRBCs) to 82 TPM (niRBCs^37°C^), further to 392.6 TPM (iRBCs^37°C^), and peaks at 1932 TPM (iRBCs^40°C^) (Figure 6, Supplementary Table 1). For other genes, such as HSPA1A, the expression with and without niRBCs is comparable, and an increase in expression is seen in the presence of iRBCs at 37°C and increases further at 40°C (TPM: 177 (without niRBCs); 160 (niRBCs); 537 (iRBCs^37°C^), 23001 (iRBCs^40°C^) (Supplementary Table 1).

**Figure 6.**
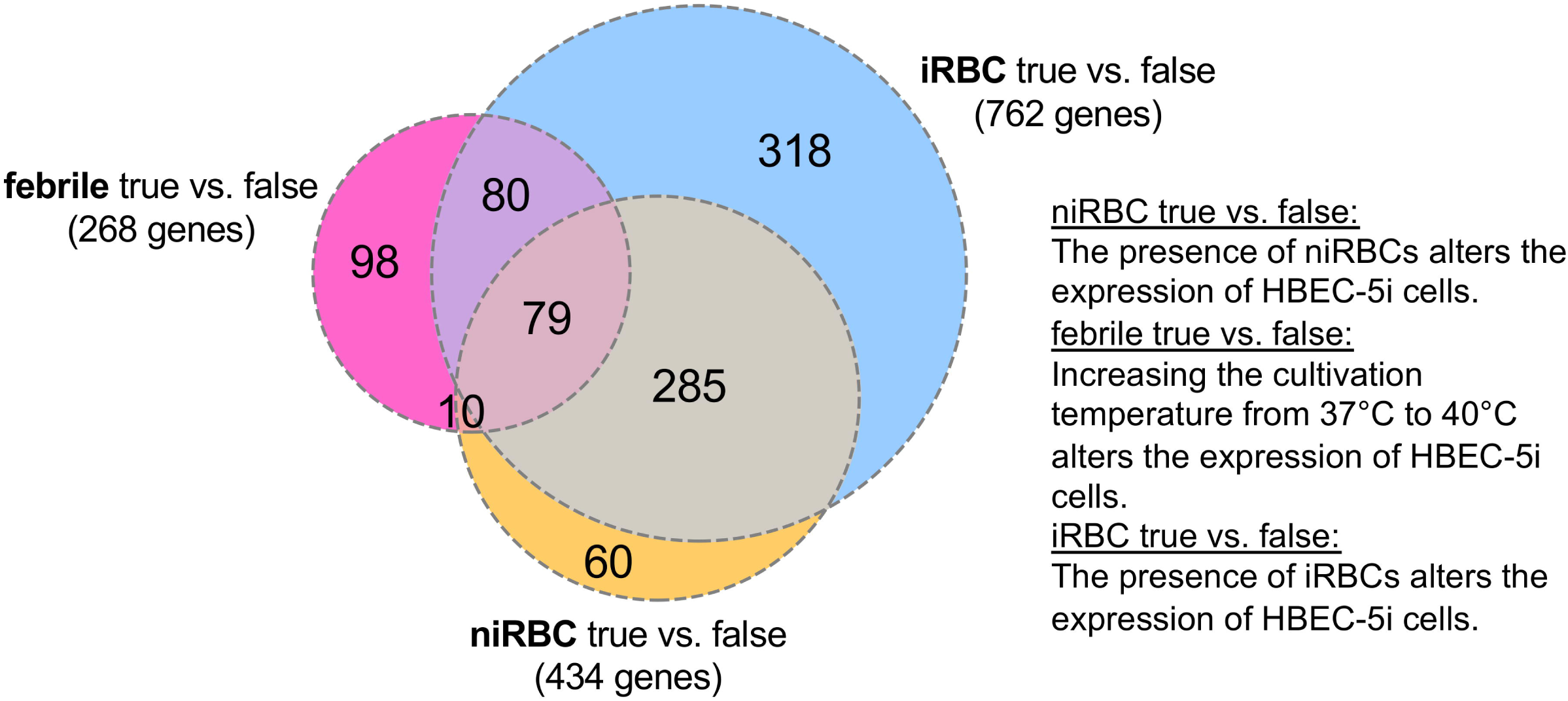
Compilation of genes whose expression is increased by the presence of iRBCs and/or an elevated temperature compared to the control. (**A**) Cytokine encoding genes. (**B**) Genes encoding metalloproteinases and NR4A nuclear hormone receptor proteins. (**C**) Genes encoding histones and histone-linker proteins, (**D**) Genes encoding proteins of the interferon-alpha/beta signaling pathway. The expression level is shown as TMP. Blue: Expression level in the presence of niRBCs^37°C^; Red: Significant increase in expression (>2-fold) due to the presence of iRBCs^37°C^; Red-brown: Significant increase in expression (>2-fold) due to the presence of iRBCs^40°C^ in comparison to iRBCs^37°C^. (**E**) Reactome analyses of the genes listed in D. The pathways to which the genes can be assigned are shown in yellow.

The g:Profiler and ShinyGO analyses of the 205 genes whose expression was up-regulated in the presence of iRBCs^40°C^ compared to iRBCs^37°C^, which were also carried out here, demonstrate that immunological signaling pathways are once again positively impacted by the combination of iRBCs and elevated temperature (Figure 4F, G). However, the analysis of the 38 down-regulated genes reveals no significantly regulated pathways.

The genes, whose expression is further increased in a temperature-dependent manner, are associated with the KEGG pathways “TNF signaling pathway” (FDR 2.96e-12), “IL-17 signaling pathway” (FDR 2.76e-11), “NF-κB signaling pathway (FDR 1.09e-09) and “cytokine-cytokine receptor interaction” (FDR 2.98e-09). Notably, for genes related to the GO-BP terms “response to cytokine” (17 genes, FDR 5.96e-19) and “protein folding” (13 genes, FDR 6.55e-23), an increase in expression is observed specifically with a temperature increase from 37°C to 40°C (Figure 5C, Supplementary Table 1).

### 2.3 Influence of iRBCs on the expression of genes encoding proteins involved in the immune response, nucleosome assembly, angiogenesis, and interferon-alpha/beta signaling pathway

The comparative analyses described above show that cytoadhesion of iRBCs leads to increased expression of genes whose encoded proteins are involved in the host’s immune response. For some of these genes, the higher temperature leads to a further increase in expression, although temperature alone has little or no effect on the expression level of these genes. The encoded cytokines can be divided into four groups. 1) Genes that show an increase in expression when co-cultured with iRBCs^37°C^, but maintain a similar level of expression when the temperature is raised to 40°C (*CXCL10, CXCL11, CCL5, CCL20, IL12A*); 2) Genes whose expression is up-regulated at 37°C and whose expression is further up-regulated when the temperature is raised to 40°C (*CXCL2, CXCL3, CXCL6, CXCL8, CCL3, CCL3L1, CCL26, IL1a, IL6*); 3) Genes whose expression increases only during co-cultivation at 40°C (*CXCL1, CXCL5, IL1b, IL32*); 4) Genes that show similar levels of expression under all conditions (*CXCL12, CXCL14, CXCL16, CCL2*) (Figure 6A, Supplementary Table 1).

Six genes whose encoded proteins can assume functions in angiogenesis are also differentially expressed. These include three genes (*MMP1*, *MMP2*, *MMP3*) encoding metalloproteinases as well as the genes (*NR4A1, NR4A2, NR4A3*) encoding the three members of the NR4A nuclear hormone receptor superfamily. For MMPs, the increased expression was only observed at elevated temperature in the presence of iRBCs. In contrast for NR4A1, NR4A2, and NR4A3, the expression increased in the presence of iRBCs and a further increase was observed after raising the temperature from 37°C to 40°C (Figure 6B, Supplementary Table 1). This suggests that the presence of iRBCs and the associated temperature increase could potentially stimulate angiogenesis, a process that is crucial for wound healing and tissue repair.

In addition, twelve genes coding for histones or histone linker proteins were significantly up-regulated in the presence of iRBCs. However, raising the temperature to 40°C did not lead to a further increase in expression; on the contrary, the expression level for four genes significantly decreased in the presence of niRBCs (Figure 6C, Supplementary Table 1).

Interestingly, an additional 14 genes were identified coding for proteins of the antiviral immune response or the interferon-alpha/beta signaling pathway (Reactome: FDR 2.22E-15) (Figures 6D, 6E, Supplementary Table 1). The presence of iRBCs alone leads to a significant up-regulation in expression. With the exception of *IL6*, there is no further increase in expression at elevated temperatures. On the contrary, for four genes (*IFI6, ISG15, MX1, MX2*) the expression level falls back to the baseline level of niRBCs (Figures 6D, 6E, Supplementary Table 1).

### 2.4 The NF-κB signaling pathway and its target genes are involved in the activation of HBEC-5i cells via cytoadhesion of iRBCs

Tripathi and colleagues showed that the pro-inflammatory NF-κB pathway is central to the regulation of the expression of HBMECs modulated by iRBC cytoadhesion (Othman et al., 2023; Tripathi et al., 2009). We therefore compared our list of up-regulated genes with the genes encoding the NF-κB subunits, the inhibitors of NF-κB, the NF-κB activation cascade and the NFκB targets. Analysis of the transcriptomes revealed an up-regulation of four out of five genes encoding NF-κB subunits (*RELB*, *REL*, *NFKB1* and *NFKB2*), only one gene encoding the NF-κB subunit *RELA* was not regulated (Table 1). Furthermore, three genes encoding NF-κB inhibitor proteins (*NFKBIA*, *NFKBIE*, *NFKBIZ*) responsible for NF-κB retention in the cytoplasm were identified (Table 1). In addition, several genes encoding members of the NF-κB activation cascade and genes encoding NF-κB target genes were up-regulated in response to iRBC cytoadhesion (Table 1). Of the 66 genes identified as up-regulated and related to the NF-κB pathway, 32 genes are expressed ≥2-fold and 12 genes are expressed ≥ 1.7-fold at 37°C compared to the control (niRBCs^37°C^). The combination of elevated temperature leads to an increase in expression of 62 genes that are expressed ≥2-fold and three genes that are expressed ≥1.7-fold. As shown above, elevated temperature alone does not increase the expression of genes involved in proteins of the NF-κB pathway or its targets (Supplementary Table 1).

**Table 1.** List of genes encoding NF-κB subunits, inhibitors of NF-κB, NF-κB activation cascade and NFκB targets, that are up-regulated after cytoadhesion of iRBCs at 37°C or 40°C in compariosn to the control (niRBCs at 37°C).

|  | Gene name | Gene symbol | Ratio |  |
| --- | --- | --- | --- | --- |
|  |  |  | niRBC<sup>37°C</sup><br>vs.<br>iRBC<sup>37°C</sup> | niRBC<sup>37°C</sup><br>vs.<br>iRBC<sup>40°C</sup> |
| **NF-κB subunits** |  |  |  |  |
| 1 | Nuclear Factor Kappa B Subunit 1 | NFKB1* | 1.5 | 2.6 |
| 2 | Nuclear Factor Kappa B Subunit 2 | NFKB2* | 1.1 | 1.9 |
| 3 | RELB Proto-Oncogene, NF-KB Subunit | RELB* | 1.4 | 2.7 |
| 4 | EL Proto-Oncogene, NF-KB Subunit | REL* | 1.8 | 3.5 |
| **NF-κB inhibitory proteins** |  |  |  |  |
| 5 | NFKB Inhibitor Alpha | NFKBIA* | 2.2 | 2.7 |
| 6 | NFKB Inhibitor Epsilon | NFKBIE* | 1.0 | 2.3 |
| 7 | NFKB Inhibitor Zeta | NFKBIZ | 2.5 | 4.1 |
| **NF-κB activation cascade** |  |  |  |  |
| 8 | BCL2 Related Protein A1 | BCL2A1 | 1.5 | 7.0 |
| 9 | Baculoviral IAP Repeat Containing 3 | BIRC3* | 3.0 | 13.5 |
| 10 | CD40 Molecule | CD40* | 0.9 | 3.1 |
| 11 | C-X-C Motif Chemokine Ligand 8 | CXCL8 | 4.8 | 25.0 |
| 12 | Intercellular Adhesion Molecule 1 | ICAM1* | 1.0 | 3.0 |
| 13 | Interleukin 1 Beta | IL1B | 0.9 | 2.4 |
| 14 | Prostaglandin-Endoperoxide Synthase 2 | PTGS2 | 3.1 | 11.4 |
| 15 | TNF Alpha Induced Protein 3 | TNFAIP3* | 4.3 | 10.3 |
| 16 | TNF Receptor Associated Factor 1 | TRAF1 | 1.1 | 2.3 |
| 17 | Vascular Cell Adhesion Molecule 1 | VCAM1 | 2.7 | 5.7 |
| **NF-κ target genes** |  |  |  |  |
| **Cytokines/Chemokines and their modulators** |  |  |  |  |
| 18 | C-C Motif Chemokine Ligand 5 | CCL5 | 2.7 | 2.2 |
| 19 | C-C Motif Chemokine Ligand 20 | CCL20* | 17.5 | 28.0 |
| 20 | C-X-C Motif Chemokine Ligand 1 | CXCL1* | 0.8 | 4.4 |
| 21 | C-X-C Motif Chemokine Ligand 3 | CXCL3* | 2.9 | 13.4 |
| 22 | C-X-C Motif Chemokine Ligand 5 | CXCL5 | 1.2 | 5.5 |
| 23 | C-X-C Motif Chemokine Ligand 6 | CXCL6* | 1.9 | 6.8 |
| 24 | C-X-C Motif Chemokine Ligand 11 | CXCL11 | 3.6 | 2.9 |
| 25 | Interleukin 1 Alpha | IL1A | 3.1 | 8.8 |
| 26 | Interleukin 6 | IL6* | 5.2 | 14.3 |
| 27 | Epstein-Barr Virus Induced 3 | EBI3* | 0.9 | 2.3 |
| 28 | LIF Interleukin 6 Family Cytokine | LIF* | 1.8 | 2.9 |
| **Immune receptors** |  |  |  |  |
| 29 | CD83 Molecule | CD83* | 0.8 | 3.6 |
| 30 | Solute Carrier Family 27 Member 3 | SLC27A3 | 1.8 | 3.0 |
| **Proteins involved in antigen presentation** |  |  |  |  |
| 31 | Complement C3 | C3 | 0.9 | 14.4 |
| **Cell adhesion molecules** |  |  |  |  |
| 32 | Neural Cell Adhesion Molecule 2 | NCAM2 | 1.8 | 1.9 |
| **Stress response genes** |  |  |  |  |
| 33 | Superoxide Dismutase 2 | SOD2* | 1.8 | 2.7 |
| 34 | MX Dynamin Like GTPase 1 | MX1 | 2.5 | 1.2 |
| **Regulators of apoptosis** |  |  |  |  |
| 35 | CD274 Molecule | CD274 | 3.8 | 7.1 |
| **Growth factors, ligands, and their modulators** |  |  |  |  |
| 36 | Inhibin Subunit Beta A | INHBA* | 1.7 | 2.2 |
| 37 | Colony Stimulating Factor 3 | CSF3 | 0.3 | 2.7 |
| 38 | Colony Stimulating Factor 2 | CSF2 | 0.8 | 46.3 |
| 38 | Secreted Phosphoprotein 1 | SPP1 | 1.7 | 2.5 |
| 40 | Vascular Endothelial Growth Factor C | VEGFC | 1.2 | 2.3 |
| **Early response genes** |  |  |  |  |
| 41 | KLF Transcription Factor 10 | KLF10 | 2.2 | 3.6 |
| **Transcription factors and their modulators** |  |  |  |  |
| 42 | Activating Transcription Factor 3 | ATF3* | 3.3 | 9.0 |
| 43 | CCAAT Enhancer Binding Protein Delta | CEBPD | 2.4 | 3.4 |
| 44 | Fos Proto-Oncogene, AP-1 Transcription Factor Subunit | FOS | 1.3 | 7.4 |
| 45 | JunB Proto-Oncogene, AP-1 Transcription Factor Subunit | JUNB* | 3.1 | 2.1 |
| 46 | MAF BZIP Transcription Factor F | MAFF* | 1.6 | 2.0 |
| 47 | Nuclear Factor, Interleukin 3 Regulated | NFIL3* | 2.6 | 4.9 |
| 48 | Nuclear Receptor Subfamily 4 Group A Member 1 | NR4A1* | 6.7 | 21.0 |
| 49 | Nuclear Receptor Subfamily 4 Group A Member 2 | NR4A2 | 25.6 | 34.6 |
| 50 | Snail Family Transcriptional Repressor 1 | SNAI1 | 3.1 | 2.9 |
| 51 | SRY-Box Transcription Factor 9 | SOX9 | 1.1 | 3.5 |
| 52 | TNF Alpha Induced Protein 6 | TNFAIP6 | 2.3 | 5.0 |
| 53 | KLF Transcription Factor 6 | KLF6* | 3.0 | 3.0 |
| 54 | Phorbol-12-Myristate-13-Acetate-Induced Protein 1 | PMAIP1 | 2.6 | 3.5 |

| **Enzymes** |  |  |  |  |
| --- | --- | --- | --- | --- |
| 55 | Dual Specificity Phosphatase 1 | DUSP1* | **2.2** | **3.9** |
| 56 | Dual Specificity Phosphatase 10 | DUSP10 | **1.7** | **5.2** |
| 57 | Indoleamine 2,3-Dioxygenase 1 | IDO1 | 1.3 | **2.3** |
| 58 | Interferon Stimulated Exonuclease Gene 20 | ISG20* | **2.3** | **1.8** |
| 59 | Matrix Metalloproteinase 1 | MMP1 | **2.9** | **4.8** |
| 60 | Matrix Metalloproteinase 3 | MMP3 | 1.1 | **8.6** |
| 61 | Pim-1 Proto-Oncogene, Serine/Threonine Kinase | PIM1 | **3.1** | **2.3** |
| 62 | Protein Phosphatase 1 Regulatory Subunit 15A | PPP1R15A* | **1.8** | **5.4** |
| 63 | Spermidine/Spermine N1-Acetyltransferase 1 | SAT1 | **1.9** | **2.0** |
| **Miscellaneous** |  |  |  |  |
| 64 | Interferon Induced Protein 44 Like | IFI44L | **4.0** | **2.0** |
| 65 | Growth Arrest And DNA Damage Inducible Alpha | GADD45A* | **2.2** | **4.5** |
| 66 | Growth Arrest And DNA Damage Inducible Beta | GADD45B | **1.8** | **3.0** |

List of genes including NF-κB subunits, inhibitors of NF-κB, the NF-κB activation cascade and NFκB targets were taken from (https://www.bu.edu/nf-kb/gene-resources/target-genes/) and from Tripathi and colleagues (Othman et al., 2023; Tripathi et al., 2009). \*Genes were also identified by Tripathi et al., 2019 as up-regulated. Genes that are upregulated $\geq 1.7$ -fold are printed in bold.

## 3 Discussion

*P. falciparum* malaria triggers a potent host immune response that plays a pivotal role in the disease’s pathogenesis through heightened systemic and local inflammation. Our comprehensive analysis, in line with previous studies, reveals that the cytoadhesion of iRBCs exerts a profound influence on the gene expression of the ECs. Notably, binding to HBEMCs primarily impacts the signaling pathways involved in the inflammatory response, apoptosis, cell-cell signaling, signal transduction, and the NF-κB activation cascade (Othman et al., 2023; Tripathi et al., 2009). However, our research goes beyond this, suggesting that cytoadhesion is not only influenced by the expression but also by the specific *Pf*EMP1 presented on the surface of the iRBCs (Othman et al., 2023). Similar effects have also been demonstrated for HUVECs. However, some studies show that the stimulation of HUVECs by cytoadhesion of iRBCs is only possible in the presence of low levels of TNF (Chakravorty et al., 2007; Tripathi et al., 2006; Viebig et al., 2005).

In our comprehensive study, we meticulously analysed the influence of iRBC cytoadhesion on HBEC-5i cells. To ensure the accuracy of our results, we used an iRBC population enriched for binding to HBEC-5i cells. This led to the creation of a homogeneous cell population presenting IT4var04 (VAR2CSA) on their surface, as CSA is the dominant antigen on HBEC-5i cells (Dorpinghaus et al., 2020). This iRBC^IT4var04^/HBEC-5i model was employed to investigate whether stimulation of HBEC-5i cells by binding of iRBCs to CSA is possible and how cytoadhesion affects the expression profile also at elevated temperatures, simulating fever.

First, the influence of the presence of niRBCs (niRBC true vs. false), iRBCs (iRBC true vs. false), and fever (febrile true vs. false) on the expression profile of HBEC-5i cells was analysed. For these true vs. false analyses, only one stimulus is considered. For example, in the case of iRBC true vs. false, all HBEC-5i transcriptomes in which iRBCs were co-incubated with HBEC-5i cells were analysed, regardless of the temperature used, and in the case of febrile true vs. false, all transcriptomes in which HBEC-5i cells were cultured at 40°C were analysed, regardless of the presence or absence of niRBCs or iRBCs. In this way, all three comparisons were analysed in a similar way. The analysis showed that the presence of niRBCs alone strongly influences on the expression profile of HBEC-5i cells. Fifty-nine genes are differentially expressed only in the presence of niRBCs. Since the presence of niRBCs is an organic component of EC homeostasis, the overlap of the two DEG analyses between niRBC (true vs. false) and iRBC (true vs. false) allows the isolation of genes that are affected by the presence of the parasite compared to those of niRBCs. However, there is a considerable overlap of 364 genes that are also differentially expressed in the presence of iRBCs. It is important to note that this analysis needs to provide more information about the magnitude of expression level changes. Particularly interesting are, for example, those genes showing up-regulated expression in the presence of niRBCs and further up-regulation in the presence of iRBCs. Some examples of genes showing this behavior are *JUNB* which encodes a subunit of the AP-1 transcription factor (TPM: 33_ no RBCs; 72_niRBCs; 227_iRBCs), *IL6* (TPM: 3_no RBCs; 13_niRBcs; 70_iRBCs), and *CXCL8* (TPM: 11_no RBCs; 82_niRBcs; 392_iRBCs). As expected, increasing the temperature from 37°C to 40°C also significantly affects on the expression profile of HBEC-5i cells, mainly affecting genes encoding heat shock proteins involved in protein folding and stress response.

*BOLA2* shows a significant increase from TPM 2.5 to 198 (almost 80-fold). This substantial up-regulation of *BOLA2*, despite its relatively unknown function, suggests a potentially crucial role in gene expression and cell signaling pathways. It has been suggested that *BOLA2* could interact with glutaredoxin-3 and act as a redox-regulated regulator of Fe-S cluster biogenesis or as a transcriptional regulator (Banci et al., 2015).

The iRBC true vs. false analysis shows that 762 genes are differentially regulated. In particular, many genes involved in immunological signaling pathways, such as the TNF signaling pathway, the IL-17 signaling pathway, the NF-κB signaling pathway, and the cytokine-cytokine receptor signaling pathway, are up-regulated. Interestingly, genes coding for proteins involved in the defense response to viruses and in nucleosome organization or assembly are expressed at higher levels.

A direct comparison of the influence of iRBCs compared to niRBCs at 37°C and 40°C shows that the positively regulated pathways are, by and large, the same as found in the iRBC true vs. false analysis.

Some of these signaling pathways have already been identified as regulated in the study by Tripathi and colleagues (Tripathi et al., 2009). In addition to the most strongly affected signaling pathways “cytokine-cytokine receptor interaction” and “cell surface receptor-linked signal transduction”, many genes of the NFκB activation cascade were also found to be positively regulated (Tripathi et al., 2009). On the other hand, some of the signaling pathways identified as regulated in the present study were not identified as regulated by Tripathi *et al*., including nucleosome assembly and the defense response to viruses (Tripathi et al., 2009).

It is known that the complex immune response elicited by *P. falciparum* infection involves the release of numerous cytokines, of which more than 30 have been detected at elevated levels in the serum or plasma of malaria patients compared to healthy individuals. These cytokines include CXCL2, CXCL4, CXCL8, CXCL9, CXCL10, CXCL11, CCL2, CCL3, CCL4, CCL5, CCL20, CCL28, IL-1b, IL-6, IL-9, IL-10, IL-12, IL-13, IL-15, IL-17A, IL-31, IL-33, IL1-RA, TNF-a, interferon-gamma (INFγ), VEGF, and G-CSF (Armah et al., 2007; Ayimba et al., 2011; Bujarbaruah et al., 2017; Che et al., 2015; Colborn et al., 2015; Herbert et al., 2015; Jain et al., 2008; Lyke et al., 2004; Prakash et al., 2006; Wilson et al., 2011). Specific cytokines (such as CXCL4, CXCL8, CXCL10, CCL2, CCL3, and CCL4) show a direct correlation with the severity of malaria infection (Armah et al., 2007; Ayimba et al., 2011; Wilson et al., 2011) and it has been shown that patients with cerebral malaria have significantly increased levels of CCL2, CCL4, CXCL4, CXCL8, CXCL10, IL-1RA, IL-6, TNF and G-CSF (Jain et al., 2008; John, Panoskaltsis-Mortari, et al., 2008; John, Park, et al., 2008; Wilson et al., 2011). Our study and previous studies, show that cytoadhesion of iRBCs increases the expression of many genes encoding cytokines (for review (Dunst et al., 2017)) even at 37°C. For nine of these 14 genes, in addition to cytoadhesion, raising the temperature from 37°C to 40°C (simulation of fever) leads to a further increase in expression of ≥ 2-fold. The latter group includes genes encoding the inflammatory chemokines CXCL2, CXCL3, CXCL6, and CXCL8, which attract neutrophils, CCL26, which attracts eosinophil granulocytes, and CCL3 and CCL3L1, which attract macrophages and other immune cells. For genes encoding the chemokines CXCL10, CXCL11, CCL5, and CCL20, there is a significant increase in expression due to cytoadhesion, but this is not further increased by the rise in temperature. These chemokines attract various immune cells, including T-cells, monocytes, and dendritic cells (for review (Hughes & Nibbs, 2018). *CXCL1* and *CXCL5* are the only chemokines encoding genes whose expression increases by the temperature shift to 40°C in the presence of iRBCs. These chemokines play a role in the attraction of neutrophil granulocytes. Recently, stimulation of ECs by plasma from malaria patients was shown to significantly increase the expression of *CXCL1* and *CXCL5* compared to healthy controls (Raacke et al., 2021). Interestingly, the increase in expression observed here for *CXCL6* and *CCL26* has not yet been described to our knowledge. *CXCL6* (granulocyte chemotactic protein-2 (GCP-2)) is one of the genes whose expression increases in the presence of iRBCs and is further increased when the temperature is raised to 40°C. As described above, CXCL6 attracts neutrophils and is a ligand of the chemokine receptors CXCR1 and CXCR2 (Fan et al., 2007). Since neutrophils are important for developing cerebral malaria, the expression of the gene was analysed in the malaria mouse model *Plasmodium berghei* ANKA (Chen et al., 2000). Still, infection of both C57BL/6 and Balb/c mice did not lead to an increase in expression in the brains of mice compared to non-infected mice. However, this was consistent with the observation that no neutrophils accumulated in the brains of mice with cerebral malaria (Van den Steen et al., 2008). To our knowledge, there is only one study that only one study has compared the plasma concentration of CCL26 in malaria patients with that of non-infected controls. It showed that the plasma concentrations of CCL26 in the malaria-exposed group were significantly lower than in the unexposed group. This observation correlates very well with our results. The TMP increases only slightly from 1 to 3 in the presence of iRBCs, only by increasing the temperature it does increase to TPM 26 (Mancebo-Perez et al., 2022). It can, therefore, be postulated that CCL26 is only important during the fever attacks of malaria. The presence of iRBCs also up-regulates the expression of genes encoding various interleukins. These include the pro-inflammatory cytokines IL-1β, IL-1α, IL-6, IL-12A, and IL-32, whereby for *IL-1a and IL6,* a more than 2-fold increase in expression is seen from co-incubation of niRBCs, to iRBC37°C to iRBC40°C. In contrast, for *IL1β* and *IL32,* an increase is only observed after co-incubation with iRBCs^40°C^. A recent meta-analysis showed that serum IL-1β levels are higher in patients with severe malaria than in patients with uncomplicated malaria. However, the level in patients with uncomplicated malaria was very similar to that of healthy controls (Mahittikorn et al., 2022). This observation fits very well with the results presented here. Only when the temperature increases from 37°C to 40°C does the expression of *IL1β* increase by about 2.5 times. *IL-32* shows the same expression profile as *IL-1β*, i.e. an increase in expression only occurs in the presence of iRBCs^40°C^. IL-32 is expressed in different isoforms in immune and non-immune cells, including ECs, and its functions include the induction of the production of several pro-inflammatory cytokines (TNF-α, IL-1β, IL-6, and CXCL8). The function of IL-32 in *P. falciparum* infection is not yet known. However, IL-32 has been described as a master regulator of intracellular bacterial, viral, and eukaryotic infections (for review (Dos Santos et al., 2018). In this context, it has been shown that in *Mycobacterium tuberculosis* (Mtb) infection, endogenous IL-32 induces the production of TNFα, IL-1β, and CXCL8, as well as the production of antimicrobial peptides and apoptosis, so that it can be assumed that IL-32 is associated with protection against Mtb infections (Bai et al., 2010; Bai et al., 2015; Montoya et al., 2014). As shown for MtB infection, the production of various cytokines such as TNFα, CXCL8, IL-1R, IFNγ, and IL-10 as well as antimicrobial peptides were induced in *Leishmania* infection (Dos Santos et al., 2018; Dos Santos et al., 2017).

Our results are very much in line with the findings of Tripathi and colleagues, who investigated the response of HBMECs to the presence of iRBCs. Significantly increased expression was detected for CXCL1, CXCL2, CXCL3, CXCL6, CXCL8, CCL20, and IL-6 (Tripathi et al., 2009). However, we were able to identify eleven further cytokines whose expression is up-regulated in the presence of iRBCs^37°C^ and/or iRBCs^40°C^. Furthermore, a number of genes encoding proteins involved in the regulation of inflammation and immune response are also increased in their expression by the cytoadhesion iRBCs compared to the control. These proteins include among others PTGS2 (COX-2) (Martin-Vazquez et al., 2023) and TXNIP (thioredoxin interacting protein) (Mohamed et al., 2021). Other genes that encode proteins involved in inflammation and immune reactions were expressed more strongly at elevated temperatures. The corresponding encoded proteins include, among others, TNFAIP3 (A20) (Wu et al., 2020), LGALS4 (galectin-4) (Cao & Guo, 2016), ATF3 (activating transcription factor 3) (Hai et al., 2010), CD74 (cluster of differentiation 74) (Beswick & Reyes, 2009), CH25H (cholesterol 25-hydroxylase) (Cho et al., 2023), and CD274 (Huang et al., 2013). Our results support observations from a number of studies that the innate immune system responds to the presence of *Plasmodium* ssp. by releasing pro-inflammatory cytokines and chemokines (Dobbs et al., 2020; Popa & Popa, 2021). The secreted cytokines/chemokines primarily play a role in controlling parasite growth and eliminating the parasites. Different regulatory cytokines can maintain the balance between pro-inflammatory and anti-inflammatory responses. However, when this balance is disrupted, the exaggerated pro-inflammatory response leads to significant adverse effects associated with severe forms of malaria and high mortality (Dobbs et al., 2020; Popa & Popa, 2021).

The three members of the NR4A nuclear hormone receptor superfamily (NR4A1 (Nur77), NR4A2 (Nur-related factor-1), NR4A3 (Neuron-Derived Orphan Receptor 1)) are also increasingly expressed in the presence of iRBCs, and a further increase was observed after elevating the temperature. In inflammatory diseases, NR4A family members are rapidly expressed and control the differentiation and activity of innate and adaptive immune cells. In addition, these proteins mediate cytokine signaling to control angiogenesis (for review (Hamers et al., 2013; Murphy & Crean, 2022)). Furthermore, three genes encoding members of matrix metalloproteinases (MMP1, MMP2, MMP3) are specifically up-regulated only in the presence of iRBCs at elevated temperatures. Among other things, MMPs are involved in the breakdown of extracellular matrix proteins and regulate functions related to inflammation, such as the activity of inflammatory cytokines and chemokines (for review (Lee & Kim, 2022)). MMP1 cleaves collagen type I and is involved in the modulation and remodeling of the extracellular matrix. MMP2 cleaves collagen type IV and is involved in angiogenesis by degrading the basement membrane of blood vessels, thus promoting the migration of endothelial cells. MMP3 can degrade various types of collagens, proteoglycans and fibronectin, and is involved in tissue remodeling and inflammatory reactions, among other things (for review (Lee & Kim, 2022). It can be argued that the induction of angiogenesis improves the oxygen supply to tissues affected by infection-induced hypoxia (Park et al., 2019). Therefore, the increased expression of genes encoding MMPs and NR4A may be a reason for the endothelial dysfunction observed in malaria.

Strikingly, 14 genes coding for proteins involved in the antiviral immune response or interferon-alpha/beta signaling pathway are up-regulated by cytoadhesive iRBCs. However, except *IL6*, there is no further increase in the expression at elevated temperatures. On the contrary, for four genes (*IFI6*, *ISG15*, *MX1*, and *MX2*), the expression level drops back to that in the presence of niRBCs. In summary, the genes up-regulated in their expression encode the pro-inflammatory cytokine IL-6, which can be induced by the presence of viruses, MX1 (MX dynamin-like GTPase 1), MX2 (MX dynamin-like GTPase 2), IFI6 (interferon alpha inducible protein 6) and ISG15 (interferon-stimulated gene 15), which inhibit viral replication, OAS2 (2’-5’-oligoadenylate synthetase 2) and OAS3 (2’-5’-oligoadenylate synthetase 3), which synthesize secondary messengers that serve to activate RNase L, which degrades viral and cellular RNA, and ISG20 (interferon stimulated exonuclease gene 20 kDa), which is an endoribonuclease that supports the degradation of viral RNA (Tanaka et al., 2014), (Deymier et al., 2022; Haller et al., 2015; Sajid et al., 2021; Sun et al., 2012; Zhu et al., 2015). In addition, the genes *BST2* (bone marrow stroma cell antigen 2), IFI27 (interferon alpha-inducible protein 27), *DDIT4*

(DNA-damage-inducible transcript 4), *GBP4* (guanylate binding protein 4), *DDX60* (DEAD-box helicase 60), which are also involved in the antiviral immune response, are up-regulated (Busse et al., 2020; Miyashita et al., 2011; Tretina et al., 2019; Ullah et al., 2021; Zhao et al., 2022). It was shown that a type I interferon (IFN) response occurs during *P. berghei* replication in the liver. A *P. berghei* RNA was identified as a pathogen-associated molecular pattern (PAMP) that can activate a type I IFN response. Thus, liver cells have sensor mechanisms that mediate a type I IFN-driven anti-parasite response (Liehl et al., 2014). However, in the case of iRBCs, it is unclear how such an response is induced in HBEC-5i cells and whether cytoadhesion is the trigger for this response or whether other factors, such as extracellular vesicles secreted by iRBCs and containing various microRNAs, might be involved (Wu et al., 2023).

Tripathi and colleagues showed that the pro-inflammatory NF-κB pathway plays a central role in regulating the expression of HBMECs after cytoadhesion of iRBCs (Tripathi et al., 2009). This observation was confirmed in our study. In addition, we showed that the combination of cytoadhesion of iRBCs and elevated temperature (40°C) affected the NF-κB pathway more than cytoadhesion of iRBCs at 37°C. At 37°C, 44 genes were up-regulated ≥ 1.7-fold, whereas at 40°C, 65 genes were up-regulated.

In conclusion, iRBCs mediated by cytoadhesion or other mechanisms such as cell-cell communication via extracellular vesicles, stimulate brain ECs. In particular, as described in other studies, several immunological pathways are activated, and the NFκB pathway seems to play a central role in the regulation of gene expression by the presence of iRBCs. An increase in the expression of genes encoding proteins involved in antiviral (-parasitic) mechanisms and angiogenesis was also observed. It was particularly striking that the cytoadhesion of iRBCs under fever conditions stimulated the expression of many genes more strongly than the cytoadhesion of iRBCs at body temperature. This was observed, for example, for various genes coding for cytokines/chemokines, MMPs, and NR4As.

## 4 Experimental Procedures

### 4.1 Cell culture

*P. falciparum* isolate IT4 (FCR3S1.2) was cultured in human 0+ erythrocytes at a hematocrit of 5% in the presence of 10% human A+ serum according to standard procedures (Trager & Jensen, 2005). Parasites were synchronized once a week with 5% sorbitol (Lambros & Vanderberg, 1979). Every two weeks, iRBCs were enriched for the presence of knobs. IRBCs were resuspended in two volumes of pre-warmed gelatin (1%, 37°C) and incubated at 37°C for 45 min (Waterkeyn et al., 2001). The supernatant containing knob-positive iRBCs was washed with RPMI 1640 medium (Gibco, Thermo Fisher Scientific, Bremen, Germany) and cultured as usual.

The HBEC-5i cell line (ATCC, CRL-3245) was cultured in DMEM/F12 medium (Gibco, Thermo Fisher Scientific, Bremen, Germany) supplemented with L-glutamine and sodium bicarbonate (2.438 g/l) at 37°C in 5% CO_2_ and passaged every 2-4 days at 70-90% confluence.

### 4.2 Enrichment of iRBCs that bind to HBEC-5i

To enrich iRBCs that bind to HBEC-5i cells, synchronized iRBCs at the trophozoite-stage (5-10% parasitemia, 1% hematocrit) were resuspended in binding medium (RPMI 1640 medium, 2% glucose, pH 7.2; Gibco, Thermo Fisher Scientific, Bremen, Germany) and co-incubated with HBEC-5i cells (70-90% confluence) for 90 min at 37°C and 5% CO_2_. After 60, 75, and 90 min, the cultures were gently shaken orbitally. This was followed by five washes with a binding medium to remove any unbound iRBCs. The remaining parasites were further cultured with 10 ml of RPMI 1640 medium and erythrocytes (5% hematocrit) to the ring-stage and harvested. Any detached HBEC-5i cells were removed with Biocoll separation solution (Merck Biochrom, Berlin, Germany). The parasites were then grown under standard conditions to a parasitemia of 5-10%, and the whole procedure was repeated seven times. Parasites enriched for binding to HBEC-5i cells were shown to express VAR2CSA (IT4_var04), which binds to CSA presented on HBEC-5i cells (Dorpinghaus et al., 2020).

### 4.3 Infected RBC-HBEC-5i co-incubation, RNA isolation, and RNAseq

To analyse the influence of cytoadhesion on HBEC-5i cell metabolism, HBEC-5i cells (70-90% confluence) were co-incubated with HBEC-5i-enriched trophozoite-stage iRBCs at a parasitemia of 5% (hematocrit 1%) in binding medium (Dorpinghaus et al., 2020). Co-incubation was performed in T-25 flasks at either 37°C or 40°C (iRBC^37°C^/iRBC^40°C^). HBEC-5i cells cultured at 37°C or 40°C +/-niRBCs (hematocrit 1%) served as controls. After seven hours of co-incubation, cells were collected in 4 ml of pre-warmed TRIzol (Thermo Fisher Scientific, Bremen, Germany), thoroughly resuspended, transferred to a 15 ml reaction tube, and incubated in a water bath at 37°C for 5 min. Samples could then be stored at -80°C or processed immediately. For RNA isolation, samples of 1 ml each were divided into 4 RNase-free reaction tubes, supplemented with 200 µl of chloroform cooled to 4°C, shaken well, and incubated for 3 min at room temperature (RT). Samples were then centrifuged at 4°C for 30 min at 12,000 *x g*, and the supernatant was transferred to a new reaction tube. RNA was precipitated by the addition of an equivalent volume of 70% ethanol. RNA was then isolated using the PureLink RNA Mini Kit (Thermo Fisher Scientific, Bremen, Germany) according to the manufacturer’s instructions. Possible contamination with genomic DNA was removed using the TURBO DNA-free kit (Fisher Scientific, Thermo Fisher, Hampton, NH, USA). RNA was then purified using the Agencourt RNAClean XP Kit (Beckham Coulter, Brea, CA, USA). Library preparation and subsequent RNA sequencing was performed on an Illumina HiSeq 4000 PE100 platform from BGI (BGI Group, Shenzhen, China).

### 4.4 Bioinformatic analyses/Statistics

The RNAseq data were mapped to the human reference genome (GRCh38.109) using CLC Genomics Workbench 21.0 (QIAGEN, Aarhus, Denmark). The raw data were submitted to NCBI-GEO (Gene Expression Omnibus) under the series record GSE266399. The Venn diagram was created using Python 3.11 and matplotlib 3.7.2. Each circle represents the differential expression (true vs. false) for each of the three parameters, niRBC, febrile and iRBC. The niRBC true vs. false group contains the genes for which the presence of niRBC affects the expression of HBEC-5i cells. The febrile true vs. false group contains genes for which an increase in temperature from 37°C to 40°C affects the expression of HBEC-5i cells. The group iRBC true vs. false contains the genes for which the presence of iRBCs alters the expression of HBEC-5i cells. Each circle represents one of the groups, with overlaps containing genes present in more than one group. Only those genes with a differential expression with a fold change of ≥ 2.0, a false discovery rate (FDR) of ≤ 0.05 and a minimum transcript per million (TPM) threshold of ≥ 2.0 were included in this analysis. The interaction network analysis was performed with the database STRING, version 12.0 (Szklarczyk et al., 2023). The following setting was used: Active interaction sources: text mining, experiments, databases, co-expression; Meaning of network edges: high confidence 0.700. Network display mode: Interactive; Network display options: Hide disconnected nodes in the network; Clustering Options: MCL clustering, inflation parameter 3; Edges between clusters: Solid lines. G:Profiler was used to perform functional enrichment analysis (https://biit.cs.ut.ee/gprofiler/gost) (Raudvere et al., 2019). The NF-κB target genes were taken from the list from the Gilmore Lab at Boston University (https://www.bu.edu/nf-kb/gene-resources/target-genes/) and from Tripathi and colleagues (Othman et al., 2023; Tripathi et al., 2009).

## Acknowledgments

The authors would like to thank Susann Ofori for excellent technical assistance. This work was supported by the Deutsche Forschungsgemeinschaft (DFG) Priority Programme “Physics of Parasitism” (SPP 2332; BR 1744/20-1 and GU 568/9-1), Leibniz Center Infection, and Jürgen Manchot Stiftung.

## Author contributions

JA: Conceptualization, Formal analysis, Investigation, Methodology, Visualization, Writing – original draft, Writing – review & editing. MB: Conceptualization, Formal analysis, Investigation, Methodology, Writing – review & editing. HT: Writing – review & editing. MPMT: Writing – review & editing. NGM: Methodology, Writing – review & editing. TR: Writing – review & editing. TG: Funding acquisition, Writing – review & editing. IB: Conceptualization, Data curation, Formal analysis, Funding acquisition, Project administration, Supervision, Visualization, Writing – original draft, Writing – review & editing.

## Data availability statement

The raw data were submitted to NCBI-GEO (Gene Expression Omnibus) under the series record GSE266399 and are available as expression browser in Supplementary Table 1.

## Abbreviated Summary

Cytoadhesion of *P. falciparum* infected red blood cells to endothelial cells leads to significant upregulation of genes associated with immune response, nucleosome assembly, NF-kappa B signaling, and angiogenesis in these cells. Increasing the temperature to 40 °C (fever) led to further upregulation of many genes, particularly those involved in cytokine production and angiogenesis.

